# Real-time Color Flow Mapping of Ultrasound Microrobots

**DOI:** 10.1101/2025.01.09.632241

**Authors:** Cornel Dillinger, Ahilan Rasaiah, Abigail Vogel, Daniel Ahmed

**Affiliations:** Acoustic Robotics Systems Lab, Department of Mechanical and Process Engineering, ETH Zurich, Switzerland

## Abstract

Visualization and tracking of microrobots in real-time pose key challenges for surgical microrobotic systems, as existing imaging modalities like MRI, CT, and X-ray are unable to monitor microscale items with real-time resolution. Ultrasound imaging-guided drug administration represents a significant advancement in this respect, offering real-time visual feedback on invasive medical procedures. However, ultrasound imaging still faces substantial inherent limitations in spatial resolution and signal attenuation, which hinder extending this method to microrobot visualization. Here, we introduce an approach for visualizing individual microrobots in real-time with Color Flow Mapping ultrasound imaging, based on acoustically induced structural oscillations of the microrobot generating a pseudo-Doppler signal. This approach enables the simultaneous localization and activation of bubble-based microrobots using two ultrasound sources operating at distinct frequency bandwidths. Our successful capture of microrobots measuring 60–80 micrometers in diameter reveals the potential of real-time ultrasonic imaging at the microscale.

**One-sentence Summary:** Advancing ultrasound imaging for real-time, microscale visualization—a critical breakthrough for therapeutic microrobots.

## Introduction

Microrobots capable of being wirelessly manipulated within the human body can revolutionize the future of medicine and offer innovative solutions in non-invasive therapeutics and surgical procedures (*1, 2*). In particular, these microscale medical assistants can enable targeted therapeutics, the delivery of drugs, genes, or cells to hard-to-reach and sensitive locations (*3*–*10*). For all their capabilities, however, a major limitation in current procedures is the absence of real-time visual feedback on the microrobot’s position in deep-seated tissues, which is crucial for their effective use in medical treatments (*11*). Various biomedical imaging modalities such as magnetic resonance imaging (MRI), X-ray and CT scanning, ultrasound imaging (US), photoacoustic imaging (PA), magnetic particle imaging (MPI), and positron emission tomography (PET), are being explored for microrobot imaging (*12*–*18*). However, each of these techniques faces its own set of challenges, such as low resolution and contrast (US), risks of ionizing radiation (X-ray, CT), the need for high-resolution real-time data (MRI), limited penetration depth (PA), and the inability to simultaneously visualize both microrobot and surrounding environment (MPI, PET). These limitations highlight the need for advanced imaging solutions capable of real-time, high-resolution visualization of microrobots within the human body (*19*).

Ultrasound imaging, with its widespread clinical use, real-time capability, and deep tissue penetration (up to 25 cm), stands as a suitable candidate for microrobot imaging (*19*). Gas-filled ultrasound contrast agents can enhance the spatial resolution of ultrasound imaging by exploiting the high acoustic contrast at gas-liquid interfaces, which provides them with high echogenicity (reflection of acoustic waves) compared to surrounding tissue (*20, 21*). This improves microscale object detectability in the commonly applied brightness mode (B-mode), in which two-dimensional grayscale images are generated from the echo intensities of in-tissue reflected ultrasound waves. Recently, magnetomotive ultrasound imaging concepts have been adapted for the real-time visualization of magnetic field-responsive microrobots. This imaging mode exploits the Doppler effect to detect shifts in ultrasound wave frequency as the waves reflect off moving objects, thereby providing information on their velocity and direction of motion. This data is then overlaid on a conventional B-mode image in the form of a colored flow map, also termed CFM-mode imaging. Current applications include discerning the motion of a magnetic nanoparticle swarm and detecting acoustic phase shifts in ultrasound signals linked to the controlled vibration of a magnetic microrobot (*22*–*24*). These studies demonstrate how CFM-mode imaging can overcome resolution and sensitivity limitations of ultrasound imaging for magnetic field-driven microrobots. Nevertheless, the efficacy of magnetomotive ultrasound imaging remains constrained by the fact that microrobots are constructed from materials having limited acoustic contrast with adjacent tissue; accordingly, the visualization has limited resolution and sensitivity, and advanced signal processing algorithms are required (*25, 26*). Recent work by Kim et al. has shown promising results in using magneto-gas vesicles to increase acoustic contrast in magnetic microagents (*27*). However, this technique has primarily been explored for tissue stiffness diagnosis, where only small agent displacements are assessed.

In this study, we present real-time CFM-mode ultrasound imaging of individual high acoustic contrast bubble-based microrobots (∼73 μm in diameter). The imaging concept relies on acoustically stimulated oscillation of the microrobots generated by encapsulated microbubbles that serve as both propulsion unit and contrast agent. Stationary but oscillating microrobots are detected by the imaging ultrasound as pseudo-moving objects, leading to random frequency shifts in the reflected ultrasound signals, referred to as pseudo-Doppler frequency shifts (*28*–*30*). Interestingly, the same effect is also observed in microrobots that are in motion. Notably, such bubble-based microrobots exhibit remarkable propulsion capabilities, driven by simple piezoelectric-driven acoustic stimulation in the liquid environment (*31*–*36*). We simultaneously harness propulsion while utilizing the strong acoustic contrast and emissions of the stimulated microrobots to enable real-time detection using a conventional ultrasound imaging system (imaging frequency *f*_US_ = 1.0 − 16.0 MHz). We demonstrate and characterize an imaging method that visualizes individual microrobots in real-time during translational motion and detects them from various angles within deep phantom tissue (up to 10 cm). Taken together, this work showcases the potential dual functionality of acoustic microrobots: wireless propulsion with real-time ultrasound imaging.

## Results

### Imaging Concept

Our demonstration of the microscale ultrasound imaging concept utilized five bubble-based acoustic microrobots arranged in a cross formation on a polydimethylsiloxane (PDMS) spin-coated glass slide. This glass slide was placed underneath an agar-based phantom block with an elliptical concavity on its bottom side, enclosing a cavity filled with deionized (DI) water. Using this setup, we can simultaneously observe single microrobots optically from below through the objective of an inverted microscope, and ultrasonically from the side using a linear array ultrasound imaging probe. In this configuration, both optical and ultrasound imaging methods visualize the plane wherein the microrobots are positioned (**Fig. 1A**).

**Fig 1.**
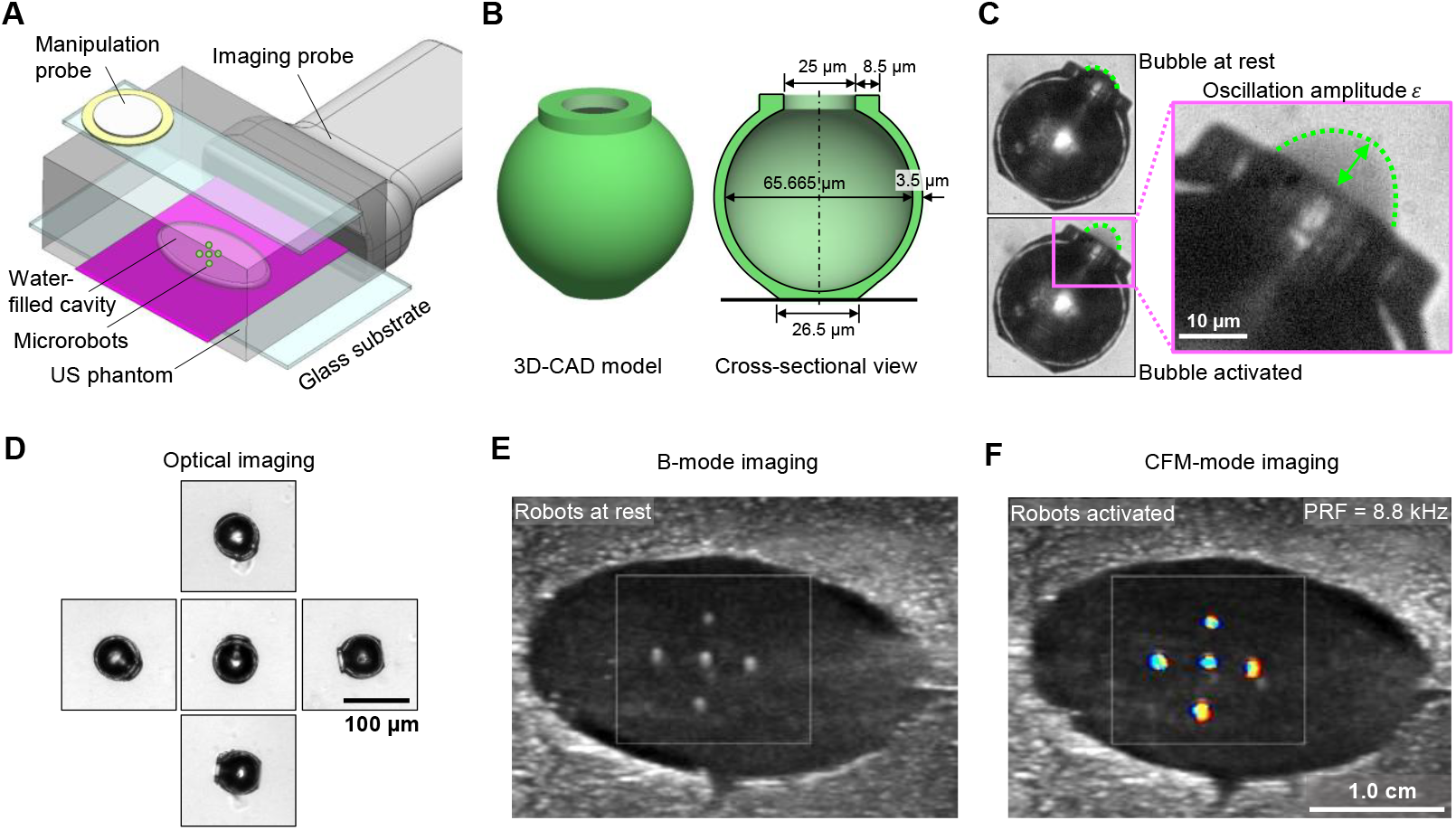
Color Flow Mapping (CFM) imaging concept. (**A**) Schematic of the experimental setup with simultaneous imaging of the same plane, optically through the glass substrate below and by ultrasound imaging from the side. (**B**) 3D-CAD model, cross-sectional view, and dimensions of the bubble-based acoustic microrobot. (**C**) When actuated with an externally applied acoustic field with frequency *f* = 100.0 kHz and amplitude V_PP_ = 7.0 V, an entrapped microbubble is stimulated to oscillate at large amplitudes. (**D**) Optical imaging of microrobots in cross formation. (**E&F**) Ultrasound imaging of microrobots using B-mode (**E**) and CFM-mode (**F**) in a conventional mobile ultrasound imaging system.

The 3D-printed bubble-based microrobots consist of hollow polymeric spheres with an outer diameter of D_outer_ ≈ 72.6 μm, a minimal wall thickness of W ≈ 3.5 μm, and a spherical orifice on one side with a diameter of A ≈ 25.0 μm (**Fig. 1B**). The hollow cavity was designed to entrap a spherical microbubble with a diameter of D_inner_ = 65.6 μm, which resonates strongly at a stimulation frequencies of *f* = 100.0 − 101.5 kHz, which coincides with the resonance frequency peaks of our custom-built manipulation probe (see also **Supplementary Information**). The manipulation probe compromises a glass slide, onto which a piezoelectric transducer disc is attached, driven by a function generator and a signal amplifier. After the 3D-printing and development process, microrobots were dried and underwent a hydrophobic surface treatment to stabilize entrapped microbubbles (see **Methods**). Subsequently, we positioned five of the microrobots in cross formation on the PDMS-coated glass slide. Due to the strong adhesion of the microrobots to the PDMS layer, the robot-containing glass slide could be flipped upside down to enclose the water-filled cavity of the ultrasound phantom (see **Methods**). After enclosing the cavity, the phantom-glass slide complex was flipped back by 180 degrees and positioned on the microscope stage for investigation. When both microrobot positioning and bubble entrapment were optically detected, we coupled the imaging (side) and manipulation (top) ultrasound probes to the phantom using ultrasonic coupling gel.

When the manipulation probe was stimulated with a sinusoidal frequency of around *f* = 95.0 − 108.0 kHz, the entrapped microbubbles exhibited strong oscillation amplitudes up to *ε* ≈ 10 μm (**Fig. 1C**). Such oscillation introduces acoustic streaming in the form of two counter-rotating vortices with a common backward jetting stream at the center of the orifice, generating propulsion in the opposite direction; a phenomenon shown in previous work (*37*–*39*). The sticky layer of PDMS on the bottom slide held the robots in position, allowing for continuous characterization and investigation of the imaging capability (**Fig. 1D**). In the water-filled elliptical chamber of the phantom, we visualized the microrobots in regular B-mode (**Fig. 1E**) and then in CFM-mode of the ultrasound imaging system (**Fig. 1F**), which provided improved contrast and visibility, consistent with earlier findings on ultrasound contrast agents (*28, 40*). In CFM-mode, the ultrasound imaging system gathers multiple signals (reflected sound waves) across a predetermined region of interest and converts this data into color tones based on the detected Doppler frequency shift, which indicates the direction and velocity of the detected object. Specifically, a blue tone denotes motion away, whereas a red tone indicates motion towards the imaging probe. In our case, it is likely that the moving object being detected is the oscillating robot-liquid interface (∼ *f* = 100 kHz), which moves back and forth with respect to the imaging probe. This can result in varying Doppler frequency-shifts generated in subsequent Doppler pulses (emitted at pulse repetition frequency (PRF), that are visualized as color-flickering CFM-mode images. In literature, it has been reported that such microbubble oscillations can lead to a loss of correlation in an imaging system’s signal analysis, resulting in so-called pseudo-Doppler signals that depict the microbubbles as random, mosaic-colored dots (*28, 29*).

Notably, when adjusting the ultrasound imaging system’s PRF (pulse values being *f*_PRF_ = 10.0, 8.8, 8.0, 7.0, 6.7, …, 1.8, 1.5, 1.2, 1.0, and 0.5 kHz while stimulating the microrobots with an acoustic field at precisely *f* = 100.0 kHz, no CFM signal was detected whenever *f*/*f*_*PRF*_ resulted in an integer. Conversely, when the stimulating frequency was not an integer multiple of the PRF (e.g., *f*_PRF_ = 8.8 kHz), the microrobots exhibited a CFM signal, observed as a red-to-blue fluctuation (**Supplementary Table S1**). This finding supports our understanding that the detecting Doppler pulse (emitted at PRF rate) and the oscillating microrobot can be in phase with each other, meaning that while the microbubble oscillates back and forth, the detecting Doppler pulse only interacts with and hence “sees” the robot’s interface as being stationary. Furthermore, this finding indicates that in our experiments the microbubbles primarily respond linearly—oscillating and generating signals—at the same acoustic frequency as their stimulation frequency, *f* = 100.0 kHz.

### Imaging Characterization

To further understand the CFM-mode imaging concept, we characterized the entrapped microbubble’s response to an acoustic frequency sweep around its main stimulation frequency bandwidth (*f* = 100.0 − 101.5 kHz). We captured the resultant microbubble actuation using both a highspeed camera and the ultrasound imaging system (**Supplementary Movie 1**). As anticipated, the microbubble’s oscillation amplitude reached its peak when the stimulating acoustic field approached the manipulation probe’s resonance frequency band *f*_*res*_ ≈ 100.0 − 101.5 kHz (**Fig. 2A**). At this frequencies, the CFM signals captured were most prominent, as shown by the highest number of colored pixels in the sonograms and in the plot of **Fig. 2B** and **2C**, indicating the correspondence of the CFM signal with the microbubble’s actuation.

**Fig 2.**
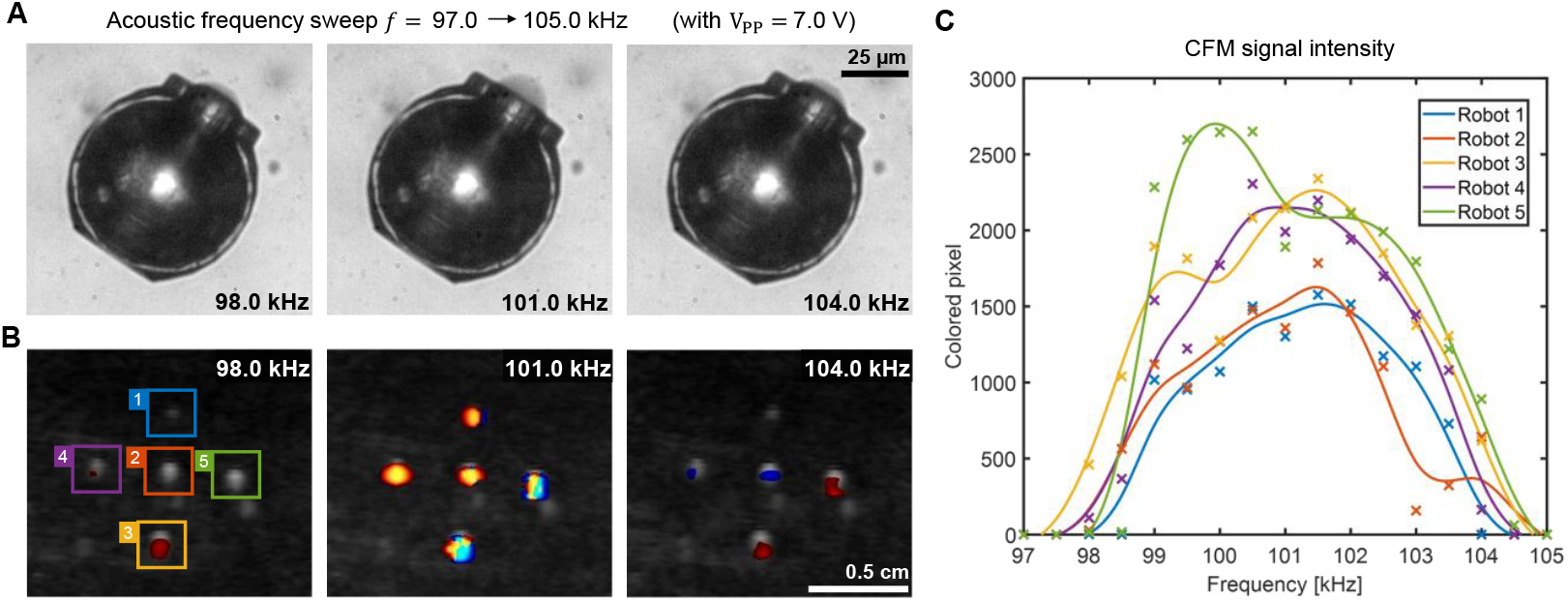
Acoustic frequency sweep. (**A**) When stimulated with acoustic frequencies in the range of *f* = 97.0 – 105.0 kHz, largest microbubble oscillation amplitudes were observed near the manipulation probe’s resonance frequency band of *f*_*res*_ ≈ 100.0 − 101.5 kHz. (**B**) Similarly, the most pronounced CFM signals were captured at *f* ≈ 100.0 − 101.5 kHz, indicated by the brightly-colored dots representing the microrobots placed in cross formation. (**C**) Plot of the average number of colored pixels in each frame of recorded CFM-mode videos versus applied frequency; imaging analysis was performed by a custom-coded pixel counter algorithm.

Next, we conducted acoustic amplitude sweep experiments. When we exposed microrobots to an acoustic field of *f* = 101.0 kHz and voltage peak-to-peak (V_PP_) amplitudes of V_PP_ = 0.7 – 14.0 V, we observed that increasing the voltage led to a larger oscillation amplitude, which was visualized in both optical and CFM-mode ultrasound imaging modalities (**Fig. 3A, 3B**, and **Supplementary Movie 2**). When plotting the total number of colored pixels in relation to applied peak-to-peak voltage amplitudes, a buckling was observed at V_pp_ ≈ 7.0 V, followed by a plateau in signal response (**Fig. 3C**). In our setup, operating microrobots in the power range V_pp_ < 40.0 V has proven sustainable for investigations spanning several minutes. Applied amplitudes exceeding V_pp_ > 50.0 V often caused microbubbles to squeeze out through the orifice or resulted in bubble collapse. In **Fig. 3D**, we showcase the importance of continuous microbubble entrapment for ultrasonic detection of the microrobots. Without any microbubble trapped, no CFM signal is generated; furthermore, in B-mode (imaging frequency *f*_US_ = 10.9 − 14.0 MHz), none of the five microrobots are distinguishable. Furthermore, to verify that a robot’s orifice orientation is not crucial for imaging, we positioned four microrobots in various orientations relative to the imaging probe (**Supplementary Fig. S1**). We detected no correlation between the color patterns in the CFM signal and the relative orientation of a microrobot’s orifice.

**Fig 3.**
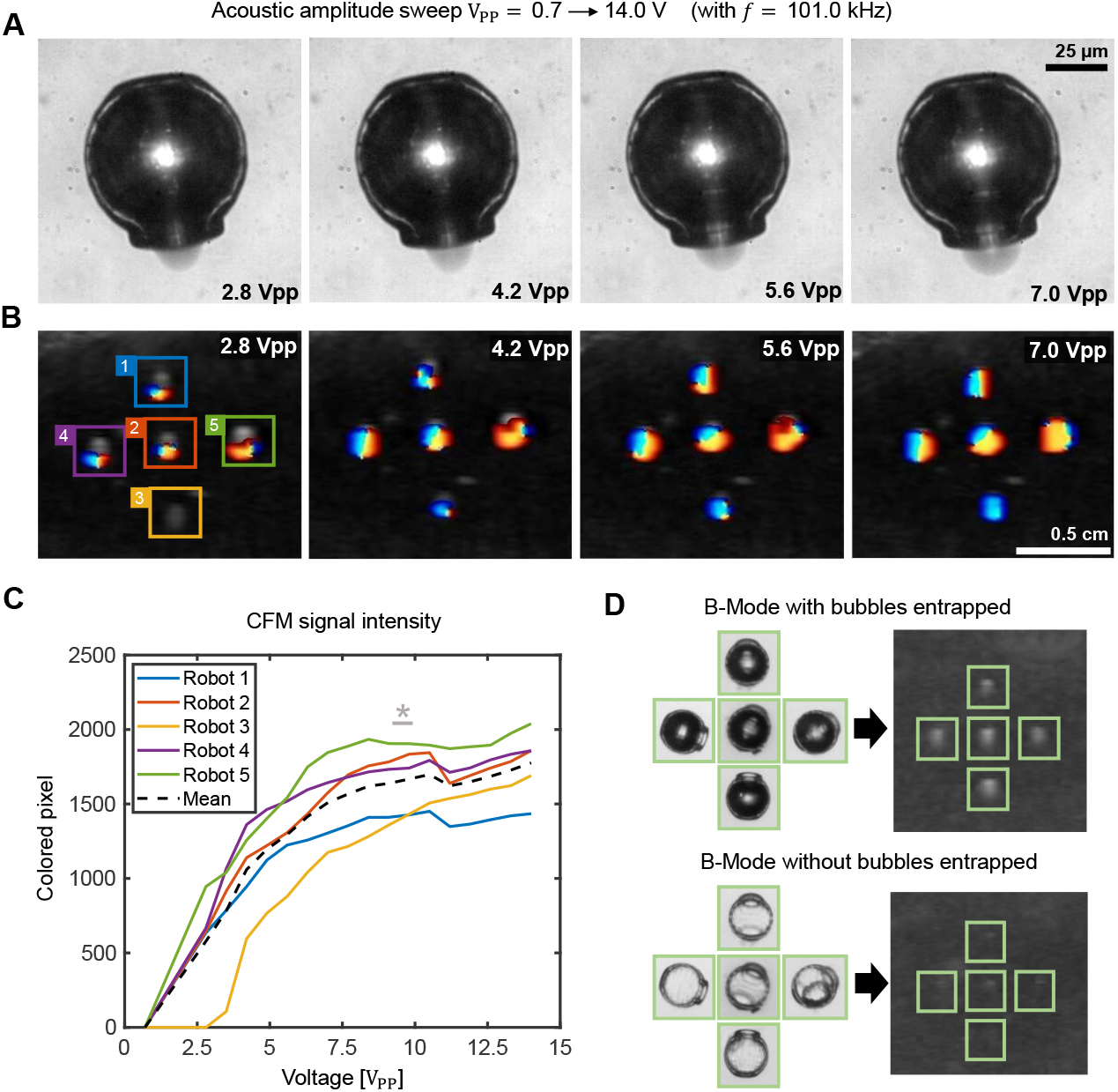
Acoustic power sweep. Microrobots exposed to increasing voltage peak-to-peak amplitudes ranging from V_PP_ = 0.7 − 14.0 V were imaged using (**A**) optical microscopy and (**B**) CFM-mode ultrasound imaging. (**C**) Plot of the average number of colored pixels in each frame of recorded CFM-mode videos versus applied power. The asterisk * indicates a set of linear interpolated (missing) data points at the value of V_PP_ = 9.1 V. (**D**) Optical and B-mode imaging of microrobots before (top) and after (bottom) application of high power amplitudes (V_PP_ > 50.0 V). Using such high amplitudes, the entrapped microbubbles often vanish, making ultrasonic detection challenging.

A vital aspect of real-time microrobot imaging within the human body is the ability to effectively detect the microrobots from any angle and over the distance required in the given situation. In this regard, ultrasound imaging benefits from its simple and portable nature, which allows medical personnel to adapt and optimize the imaging procedure according to need. In **Fig. 4**, we demonstrate the detection of our bubble-based acoustic microrobots from various imaging angles. For this experiment, we designed a tissue phantom with a curved surface that allowed us to assess the imaging probe’s ability to visualize microrobots over a 180° perspective. Inside the water-filled chamber of the phantom, five microrobots were positioned in two rows with two in one row and three in the other. As in previous experiments, the simultaneous visualization of all five microrobots was facilitated by initially holding the imaging probe in a parallel plane to the glass substrate at angles θ_Probe_ = 0° and 180° (**Fig. 4A**). We then moved the imaging probe upward toward imaging angles of 30°, 60°, and 90°, during which the microrobots remained detectable, exhibiting consistent signal intensity. We attribute this phenomenon to the radial oscillation patterns of the activated microbubbles entrapped within the microrobots. Further manual (hand-held) guidance of the imaging probe demonstrated that the oscillation signal could be detected from any imaging angle (**Fig. 4B**). However, when the imaging plane, indicated in purple, deviated from the planes at θ_Probe_ = 0° and 180°, simultaneous visualization of all five microrobots became unattainable. This is not surprising as the imaging ultrasonic beam has limited height and depicts only objects within its imaging plane. When the imaging angle is adjusted away from the parallel-to-ground plane, the imaging plane interferes only with one of the rows of microrobots. In **Fig. 4C**, we demonstrate at θ_Probe_ = 60°, it is feasible to adjust the imaging plane to display both rows; however, imaging both rows with full intensity at the same time remains unachievable (see **Supplementary Movie 3**). This result highlights the challenge of locating the correct imaging plane when the objects to be detected are of microscale dimensions.

**Fig 4.**
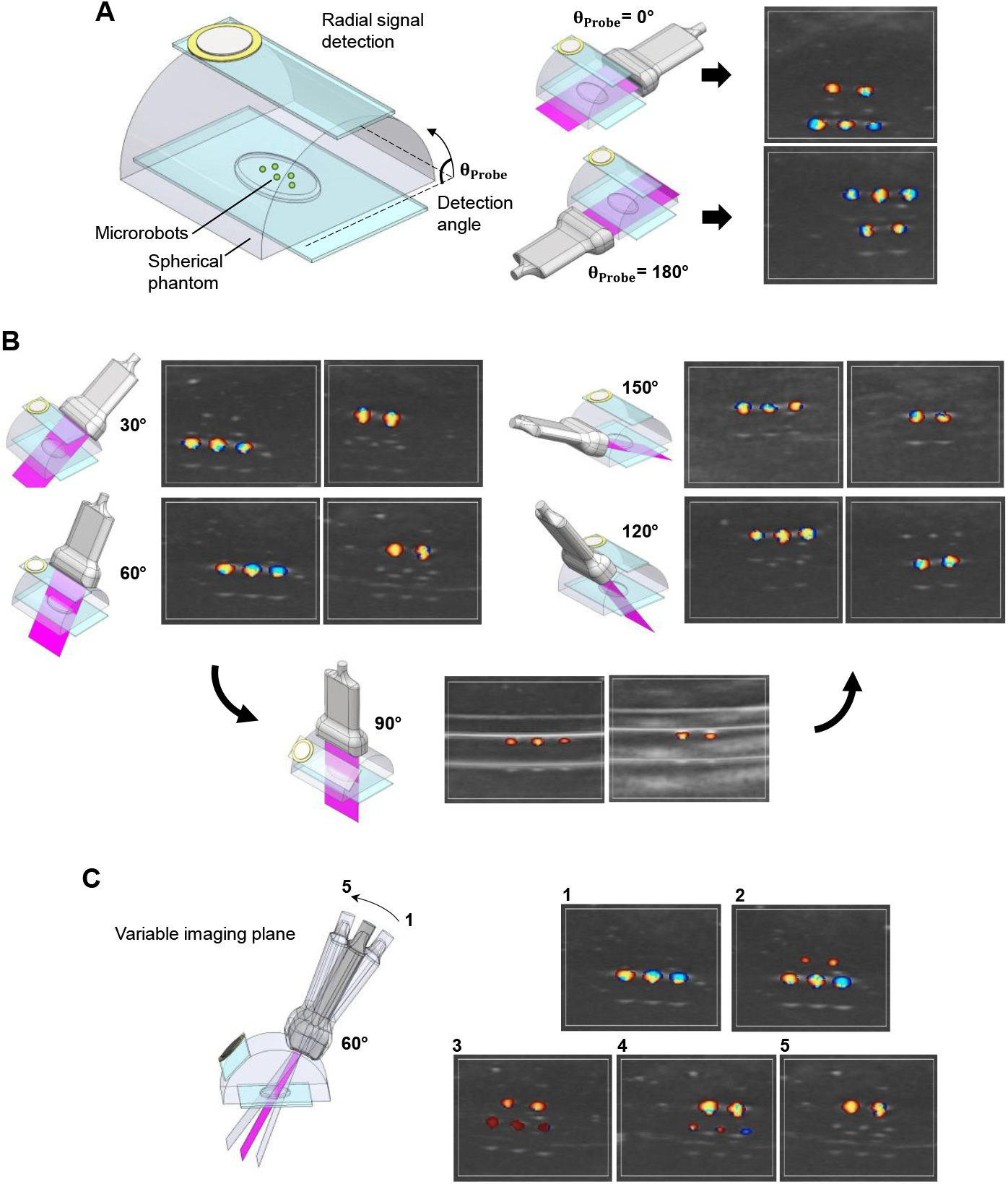
Radial signal detection. (**A**) Schematic showing the placement of five microrobots when analyzing microrobot detectability from various imaging angles. Both rows of microrobots could be imaged in plane with the glass substrate when the ultrasound imaging probe was held at θ_Probe_ = 0° or 180°. (**B**) Successful imaging of microrobots at imaging angles ranging from θ_Probe_ = 30° − 150°. In these cases, the surface of the glass substrate does not align with the ultrasound imaging plane. Therefore, the two rows of microrobots were separately visualized by partially rotating the imaging probe around the point of contact with the ultrasound phantom, as illustrated in (**C**).

### Deep Tissue Visualization

Real-time high-resolution visualization of microrobotic therapeutics in deep-seated tissue remains a challenging task. In **Fig. 5**, we present real-time CFM-based ultrasound imaging of our microrobots in agar-based tissue phantoms, covering depths up to D = 10 cm. First, we validated presence of five microrobots in cross formation from an imaging distance of about D = 3 cm (**Fig. 5A**). After confirming the presence of the trapped microbubbles under the microscope, we repositioned the imaging probe to the opposite side of the tissue phantom and began detecting signals from the microrobots at distances as far as D = 10 cm. Scanning the DI-water-filled chamber to detect the microrobots includes the consideration of the ultrasound beam’s refraction at the tissue-water interface according to Snell’s law (*41*). When ultrasound passes through an interface between two media with varying propagation speeds, the sound beam, i.e., the imaging plane, becomes bent leading to a non-intuitive imaging angle of the probe required to find the desired imaging plane. The usage of the CFM imaging mode helped to identify the microrobots in deep tissue situations as illustrated in **Fig. 5B**. When the acoustic field was stimulated at *f* = 101.0 kHz and V_pp_ = 21.0 V, a clear visible signal of each microrobot was detected. However, in the absence of an acoustic stimulus, the location of the microrobots was hard to identify since only a weak, greyish-blurred region indicated their presence (**Supplementary Movie 4**). This result emphasizes the benefits of using CFM-mode in addition to conventional B-mode imaging when detecting microscale objects equipped with microbubbles.

**Fig 5.**
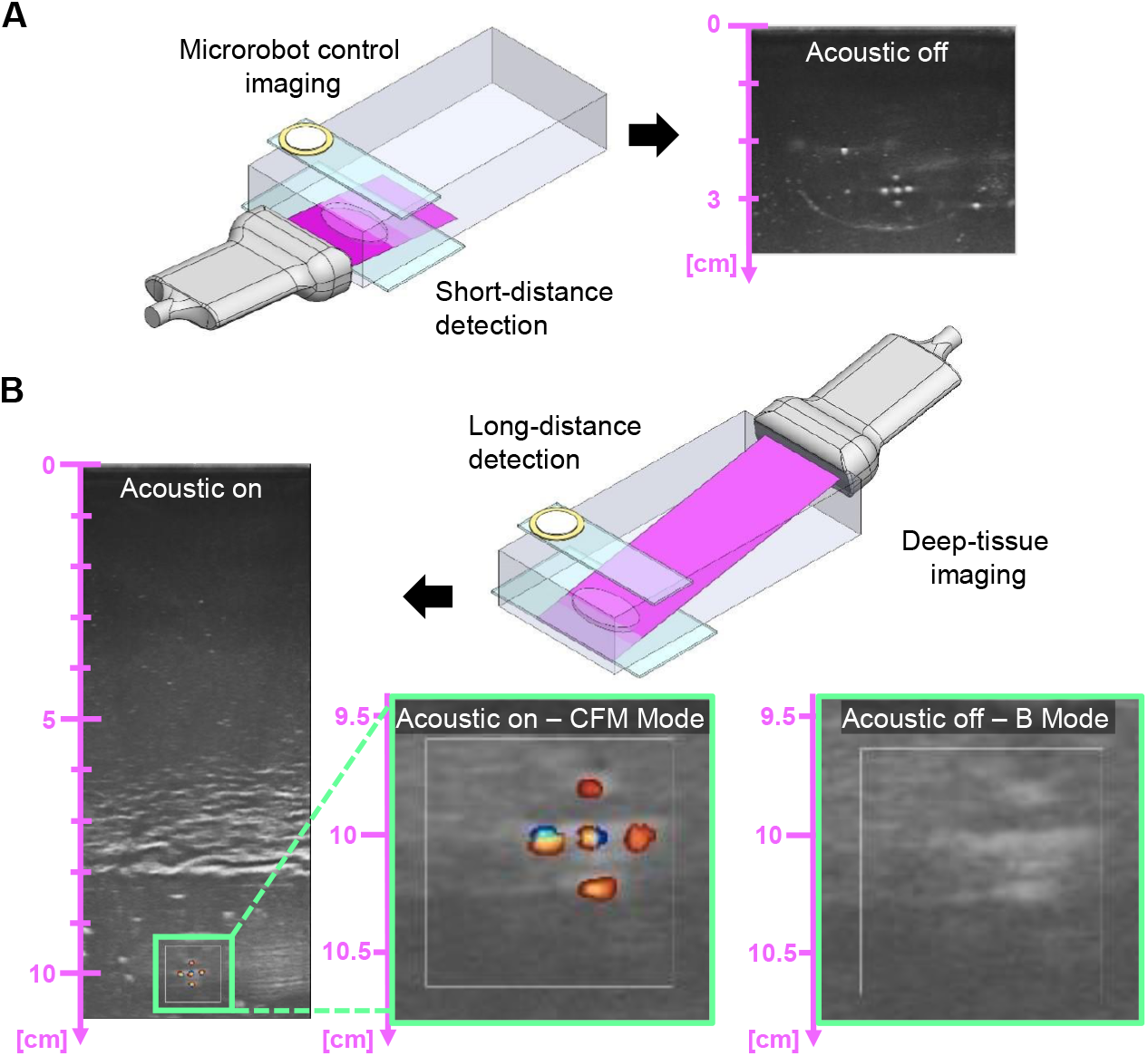
Deep tissue visualization. An elongated ultrasound tissue phantom was used to test microrobot detectability at different depths. (**A**) Visualization in B-mode at the short depth of approximately D = 3 cm. (**B**) Imaging of microrobots from the opposite side, over a distance of approximately D = 10 cm. In the absence of acoustic stimulation (B-mode), detecting the microrobots is challenging. However, when acoustics are enabled, the CFM signal clearly detects all five microrobots.

### Real-time Motion Imaging

In this section, we aim to emphasize one of the biggest advantages of ultrasound imaging – the real-time representation of live data. For the design of precise microrobotic-assisted therapeutics such as the application in targeted drug, gene, or cell delivery, microsurgery, or diagnostic sensing continuous real-time feedback on the microrobots’ performance is essential. This involves monitoring the microrobots’ movement towards the designated site, verifying the successful execution of their task, and confirming that the treatment has not exacerbated the situation. In this context, we showcase the successful real-time detection of our acoustically driven bubble-based microrobots in translational motion. To enable free movement of the microrobots in this experiment, we replaced the sticky PDMS spin-coated glass substrate with a non-coated smooth glass slide. When a bubble-based microrobot is activated on such a smooth substrate and immersed in an Newtonian fluid (DI-water), it starts to generate acoustic streaming and reorients itself, aligning its orifice perpendicular to the substrate before undergoing spherical motion patterns (**Fig. 6A**)(*38*). A secondary effect, supported by the smooth glass substrate, resulted in the microrobots’ immediate movement toward the cavity wall as soon as the imaging probe was positioned on the tissue phantom. Upon entering the DI-water-filled chamber, the ultrasonic waves from the imaging probe induced acoustic bulk streaming (see also **Supplementary Figure S2**), known in the literature as Eckart streaming (*42*). In prior experiments, this effect remained negligible because of the substantial adherence generated by the PDMS-coated glass substrate, which held the microrobots in position. Nevertheless, the acoustic bulk streaming, flowing in direction away from the imaging probe, resulted in the microrobots being pushed towards the wall of the cavity situated on the opposite side from imaging probe. This represents the initial position of the microrobot depicted in the first frame of **Fig. 6B**. We then applied an acoustic stimulation of *f* = 101.0 kHz and V_pp_ = 8.4 V on the manipulation probe to actuate the microrobot and set it into motion. This effective wireless acoustic propulsion principle has been demonstrated in various work (*32, 33*). In our specific scenario, the propulsive force facilitated the microrobot’s detachment from the wall, leading to a characteristic trochoidal motion patterns (*38*). After completing two spherical trajectories, the microrobot returned to the wall where the acoustic stimulation was turned off. We simultaneously recorded the microrobot’s motion with a CCD camera and the ultrasound imaging system in CFM-mode, therefore demonstrating the capability to real-time image the motion of bubble-based microrobots (**Fig. 6B** and **Supplementary Movie 5 & 6**). In an additional experiment, using 6-µm flow tracer immersed in the robots’ aquatic environment, we verified the situation where stationary microrobots are exposed to a background flow, i.e. the previously introduced steady-state bulk streaming. Due to acoustic waves reflected from the 6-µm flow tracers, the CFM-mode correctly detected a steady-streaming, however, it did not prevent the visualization of the microrobots (**Supplementary Figure S2**). In conclusion, we realize that the operating imaging probe can introduce streaming effects into the aquatic environment of the microrobots. However, due to the powerful bubble-based propulsion employed, the microrobots can overcome this environmental stress and remain visible, even in the presence of background flow.

**Fig 6.**
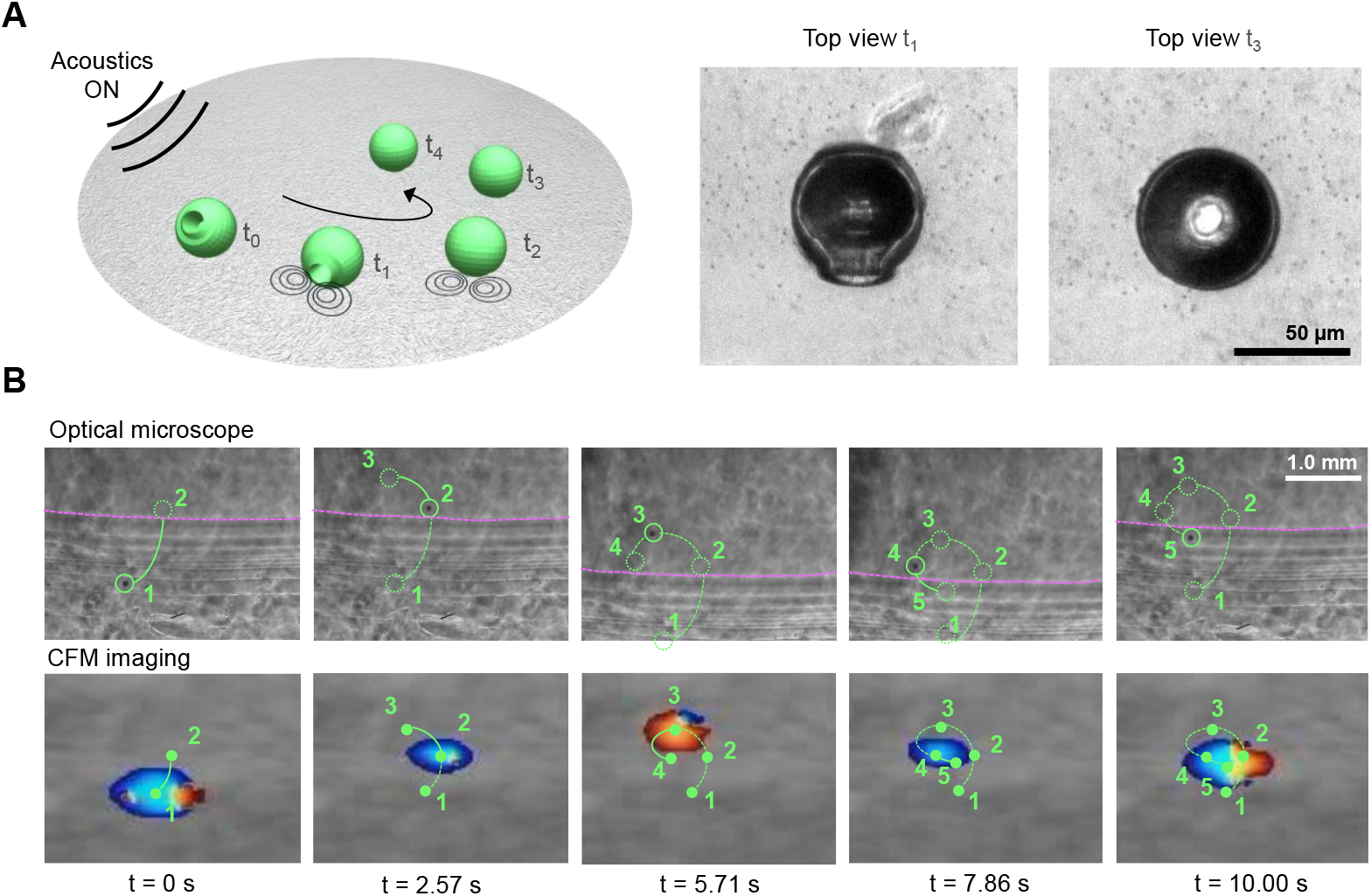
Real-time motion imaging. (**A**) When acoustics is activated, the bubble-based microrobot starts to generate acoustic streaming and reorients itself, aligning its orifice perpendicular to the substrate before undergoing spherical motion patterns. (**B**) Image sequence depicting simultaneous capturing of a swimming microrobot under an optical microscope and ultrasound imaging in CFM-mode. Acoustic stimulation with *f* = 101.0 kHz and V_pp_ = 8.4 V resulted in translational motion along a spherical trajectory recorded by both imaging modalities.

## Discussion

Using the presented imaging method for acoustic bubble-based microrobots, the inherent advantages of ultrasound imaging — such as cost-effective and portable equipment, real-time data representation, and deep tissue penetration — are enhanced by the effective visualization of microobjects equipped with high acoustic contrast. Entrapped microbubbles simultaneously act as the propulsion unit driving the microrobots, and as a contrast agent enhancing their detectability. The imaging concept leverages the improved sensitivity of the CFM-mode in conventional ultrasound imaging systems to real-time visualize stationary and moving microobjects, e.g. microrobots. In previous work, these types of microrobots demonstrated impressive swimming speeds of up to 350 mm/s (≈ 17,500 body lengths per second), making them well-suited to withstand the high shear stress levels experienced by microscale robots (*32*). While challenges remain, such as the temporal bubble instability caused by rectified diffusion, recent advancements have significantly improved bubble stability, enabling these microrobots to navigate physiologically relevant fluids, achieve 3D steerability, and even manipulate single particles (*4, 33*).

In addition to real-time imaging capabilities, we briefly demonstrate in **Fig. 7** how a potential drug delivery task could be visualized using our proposed CFM-based imaging approach. In **Fig. 7A**, we illustrate how a microrobot orients its orifice and is attracted toward the cavity’s wall governed by the secondary Bjerknes force effect, introduced by oscillating bubbles near rigid walls (*43*). A bubble-based microrobot can be manipulated to adhere to the wall and release from it once the acoustics are turned off (**Fig. 7B** and **7C**). This controllable adherence to the wall is accompanied by acoustic microbubble streaming toward the wall (**Fig. 7B** and **Supplementary Movie 7**), which can be used to direct dissolved drugs towards targeted regions in the cavity’s wall. Furthermore, seeing the resultant CFM-Mode signal of such an event (**Fig. 7C**), we envision that our real-time visualized microrobots can accomplish targeted tasks such as exerting stress (vibration) to specific locations in order to trigger mechanotransducive mechanisms.

**Fig 7.**
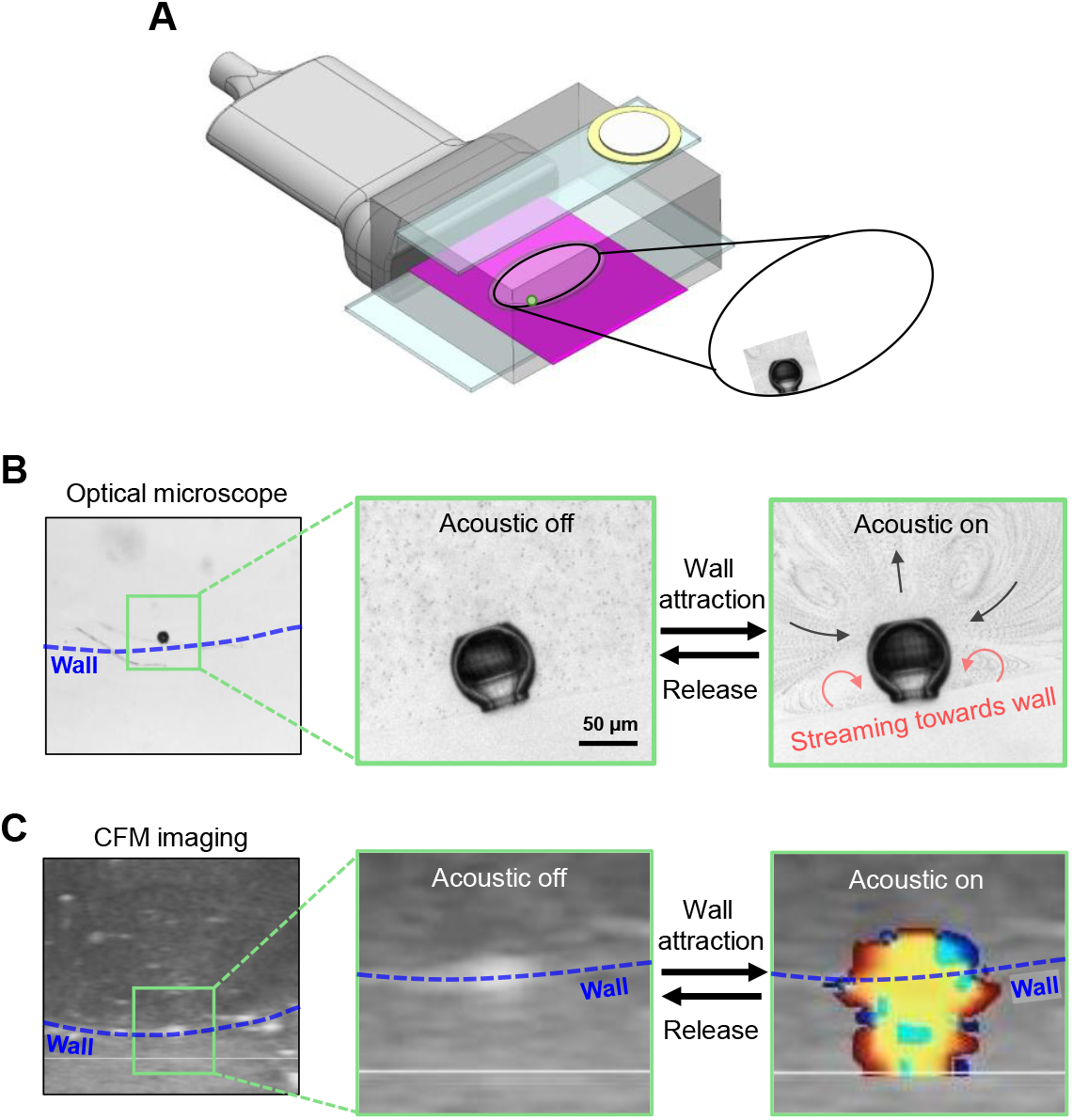
Real-time ultrasound imaging feedback in a drug delivery approach. (A) Schematic illustrating direction of a microrobot toward the wall of the cavity in the ultrasound tissue phantom. (B) Upon reaching the wall, the microrobot orients its orifice towards the wall, introduced by acoustic stimulation with *f* = 103.4 kHz and V_pp_ = 35.0 V. Activating and deactivating the acoustics controls the microrobot’s attraction to and release from the wall, accompanied by acoustic streaming directed towards the wall. (C) CFM imaging reveals the microrobot’s actuation at the wall when acoustics is turned on.

Advancing real-time feedback-guided microrobotics promotes its widespread clinical adoption towards biomedical applications (*44*). Here, these applications could include the treatment of bladder cancer through localized drug delivery, addressing stomach ulcers by precisely targeting ulcerated areas with therapeutic agents, and managing inflammatory bowel disease, such as Crohn’s disease or ulcerative colitis, by delivering anti-inflammatory drugs directly to affected intestinal tissues (*45*). However, these advancements not only can increase treatment precision but also broaden the applications of microrobotics in healthcare. Our findings have the potential to influence future research and innovation in diagnostic disease mapping, localized neuromodulation, and real-time tissue monitoring. Ultimately, this study reports a method by which ultrasound imaging can overcome its resolution deficit at the microscale and enable effective real-time observation and guidance of next generation microrobotic therapeutics and diagnostics.

## Materials and Methods

### Microrobot Fabrication

The bubble-based microrobots were 3D printed using a two-photon polymerization system (Photonic Professional GT, Nanoscribe GmbH, Karlsruhe, Germany) with a 25x objective (numerical aperture = 0.8). We printed arrays of microrobots on indium tin oxide (ITO) – coated glass substrates using the biocompatible IP-S photoresin (Nanoscribe GmbH, Karlsruhe, Germany), enabling high feature size-resolution printing. After 3D printing, the microrobots were developed in propylene glycol monomethyl ether acetate (PGMEA, Sigma-Aldrich) for 20 min and rinsed in isopropyl alcohol (IPA) for another 5 min. To ensure hydrophobic cavities, we dried the microrobots for 20 min at 85ºC and then performed a silanization (1H, 1H,2H, 2H perfluorooctyl-trichlorosilane, Sigma-Aldrich) process in a vacuum chamber for 45-60 min.

### Ultrasound Phantom Preparation

To prepare agar gel, 8 g of agar (Agar powder, Migros, Switzerland) is mixed with 1.5 dl boiling water while ensuring continuous stirring. This stirring process is sustained for 3 mins, to ensure proper solidification of the agar gel. Subsequently, well-mixed agar solution is carefully poured into the 3D-printed mold, which has been crafted using a 3D filament printer (ENDER 3 V2). After molding, the agar gel is cooled down in a refrigerator at 5°C for a minimum of one hour.

### Experimental Setup

The setup is assembled as follows: First, microrobots are positioned in cross formation on a PDMS spin-coated (duration = 1 min, rpm = 1000 min^−1^) glass slide (38 × 75 × 1 mm) using an optical fiber (AFS 100/110/130T, Fiberguide industries, Inc.), under careful attention on an optical inverted microscope. Next, the agar phantom, prepared previously, is gently released from the mold, ensuring the chamber faces upwards. The upside-down lying chamber is then filled with DI-water. The glass slide, bearing the microrobots facing downwards, is slowly lowered over the chamber from one side to the other, with a slow approach to reduce the risk of microrobots being flushed away and to prevent air bubble formation between the chamber and the glass slide. Subsequently, the phantom with the glass slide is carefully flipped by 180º so that the phantom is now positioned on the glass substrate. To ensure optimal coupling between the manipulation probe, comprising a piezoelectric transducer (Murata, 7BB-27-4L0) attached to a conventional microscope glass slide (25 × 75 × 1 mm), and the phantom, as well as between the imaging probe and the phantom, ultrasound coupling gel (K-Y gel, medical sterile) is applied in both cases. By mounting this assembly on an inverted microscope, with optical access from below and ultrasound imaging from the side, optical verification of what is observed under ultrasound is enabled.

### Imaging and Data Analysis

For experiments, a Zeiss Axiovert 200 M inverted microscope was utilized, equipped with a high-speed camera (CHRONOS 1.4, Kron Technologies). Recorded data was analyzed using software such as ImageJ and MATLAB. The ultrasound imaging system employed in all experiments is a portable Color Doppler ultrasound scanner (Sonoscape E2) equipped with a linear array imaging probe (Sonoscape, L741).

## Author contributions

D.A. and C.D. conceived the project idea. Under the guidance of C.D., A.R. and A.V. contributed to the experimental work. C.D. was leading the article writing process. D.A. and C.D. developed the theoretical background and analysis. All authors contributed to the scientific discussion.

## Data and materials availability

All data needed to evaluate the conclusions in the paper are present in the paper and/or the Supplementary Materials.

## Supplementary Materials

Please see separate file.

