## Supplementary Information for "Real-time Color Flow Mapping of Ultrasound Microrobots"

**Affiliations**

**This PDF file includes:**

Supplementary Information
Figs. S1 to S2
Table S1
Description of Movies S1 to S7

**Other Supplementary Materials for this manuscript include the following:**

Movies S1 to S7

**Supplementary Information**

<u>Free bubble resonance frequency:</u>
The natural resonance frequency of a free microbubble under adiabatic conditions, neglecting surface tension effects, and matching the size of the microrobot's cavity in this study, can be computed as (*46*),

$$f_{res} = \frac{1}{2\pi r_0}\sqrt{\frac{3\gamma p_0}{\rho}} \approx 100.0 \text{ kHz},$$

where $r_0 \approx 32.8325$ µm represents the radius of the microbubble and $\gamma = 1.4$, $p_0 = 1\, atm$, and $\rho = 1000.0\, \frac{kg}{m^3}$ are the adiabatic index, the atmospheric pressure, and the density of the surrounding fluid.

**Supplementary Figures & Tables**

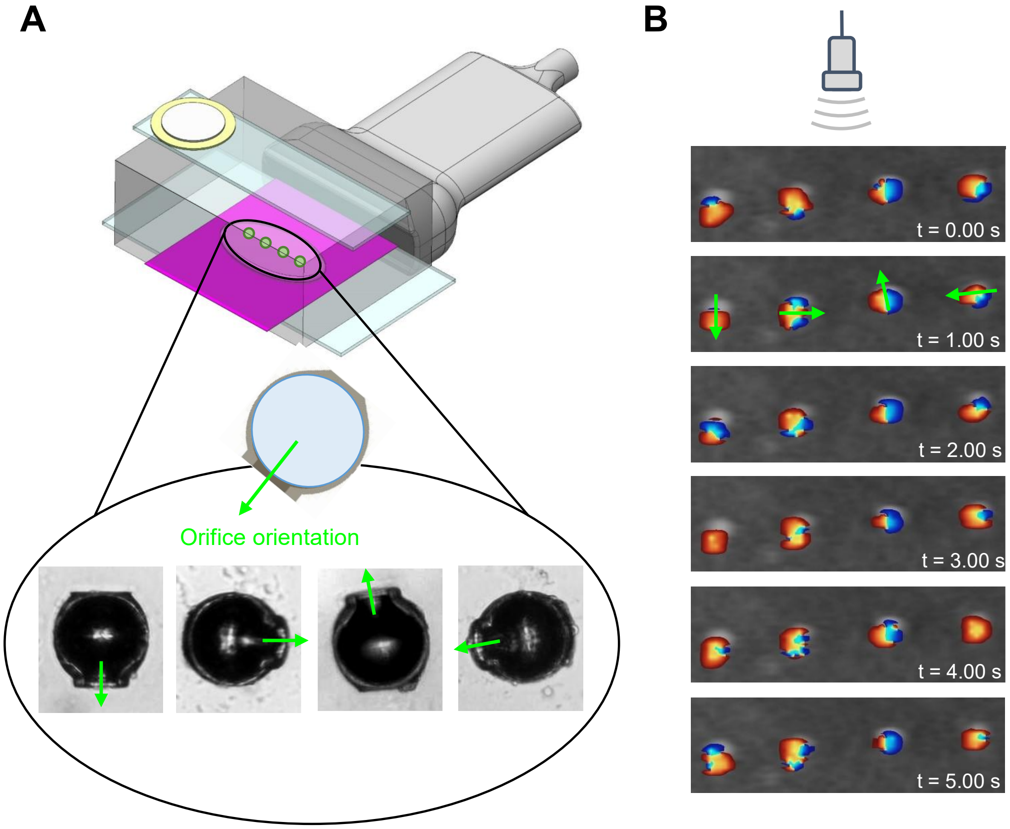


**Fig. S1. Microrobot orientation independency for detection.**
(**A**) Four microrobots positioned with different orifice orientations were placed in the ultrasound phantom. (**B**) Under stimulation with an acoustic field of $f = 103.6$ kHz and $V_{pp} = 42.0$ V, no characteristic orientation-dependent signal was detected in CFM-mode ultrasound imaging.

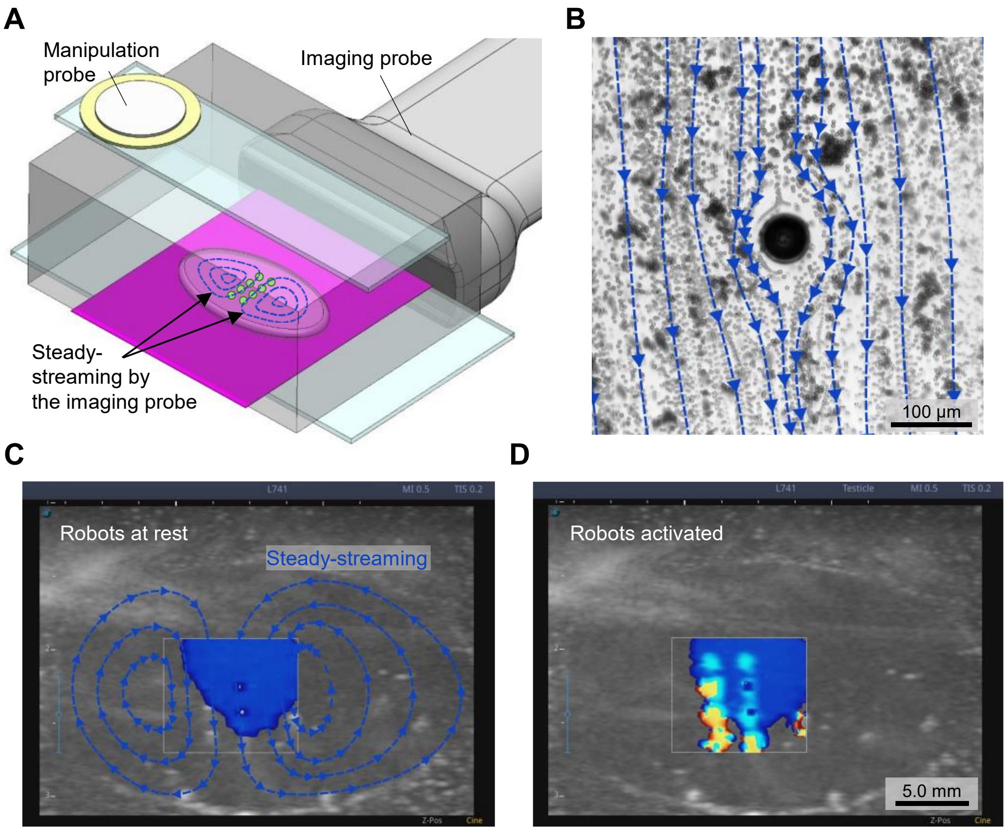


**Fig. S2. Visualization of microrobots under Doppler-detected steady-streaming.**
(**A**) Schematic of steady-streaming, i.e., Eckart streaming (*42*), within the elliptical chamber of the ultrasound phantom, generated by the imaging probe. To visualize the steady-streaming, 6 µm flow tracer particles were added to the DI-water solution. In the optical Z-stack image of (**B**), the flow field around one of the eight positioned microrobots is depicted. (**C**) Due to the flow tracers immersed in the DI-water, CFM-mode ultrasound imaging ($f_{\mathrm{PRF}} = 1.2\ \mathrm{kHz}$) of the microrobots at rest resulted in Doppler-detected steady-streaming covering the microrobots. (**D**) Activated by the manipulation probe ($f = 100.0\ \mathrm{kHz}, \mathrm{V_{pp}} = 3.5\ \mathrm{V}$), the microrobots became clearly visible using CFM-mode imaging, even amidst the Doppler-detected steady streaming.

| Acoustic field $f$ [kHz] | 100.0 | 100.0 | 100.0 | 100.0 | 100.0 | 100.0 | 100.0 | 100.0 | 100.0 | 100.0 |
|---|---|---|---|---|---|---|---|---|---|---|
| Pulse Repetition Frequency $f_{PRF}$ [kHz] | 10.0 | 8.8 | 8.0 | 7.0 | 6.7 | 6.0 | 5.7 | 5.0 | 4.4 | 4.0 |
| $\frac{f}{f_{PRF}}$ | 10.0 | 11.364 | 12.5 | 14.286 | 14.925 | 16.667 | 17.544 | 20.0 | 22.727 | 25.0 |
| CFM signal | 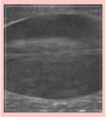 | 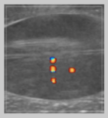 | 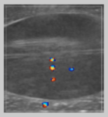 | 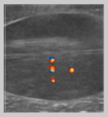 | 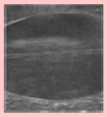 | 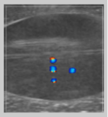 | 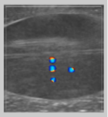 | 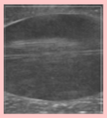 | 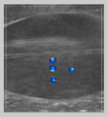 | 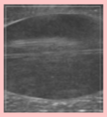 |

| Acoustic field $f$ [kHz] | 100.0 | 100.0 | 100.0 | 100.0 | 100.0 | 100.0 | 100.0 | 100.0 | 100.0 | 100.0 |
|---|---|---|---|---|---|---|---|---|---|---|
| Pulse Repetition Frequency $f_{PRF}$ [kHz] | 3.2 | 3.0 | 2.5 | 2.0 | 1.8 | 1.5 | 1.2 | 1.0 | 0.8 | 0.5 |
| $\frac{f}{f_{PRF}}$ | 31.25 | 33.333 | 40.0 | 50.0 | 55.556 | 66.667 | 83.334 | 100.0 | 125.0 | 200.0 |
| CFM signal | 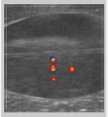 | 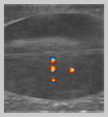 | 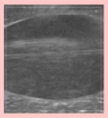 | 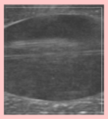 | 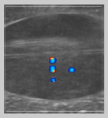 | 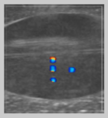 | 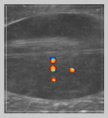 | 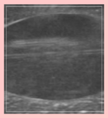 | 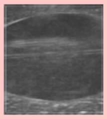 | 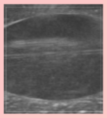 |

**Table S1. Dependence of CFM signal on the interference between the stimulating acoustic field and Pulse Repetition Frequency (PRF).**

When the stimulation frequency of the acoustic field divided by the Pulse Repetition Frequency ($\frac{f}{f_{PRF}}$), in the ultrasound imaging system resulted in an integer (except when $f_{PRF}$ = 6.7 kHz), the CFM-mode failed to detect the activated, oscillating microrobots. In this experiment, the robot positioned to the left within the cross formation remained non-responsive (no microbubble entrapped).

**Movie S1.**
Frequency sweep experiment: The response of a microbubble entrapped within a microrobot to an acoustic frequency sweep over the range of $f = 97.0 - 105.0$ kHz ($V_{PP} = 7.0$ V) is captured using a CCD camera and an ultrasound imaging system operating in CFM-mode.

**Movie S2.**
Power sweep experiment: Microrobots exposed to an acoustic field at $f = 101.0$ kHz with voltage peak-to-peak ($V_{PP}$) power amplitudes ranging from $V_{PP} = 0.7 - 14.0$ V are captured using a CCD camera and an ultrasound imaging system operating in CFM-mode.

**Movie S3.**
Variable imaging plane: At $\theta_{Probe} = 60°$, with improved proficiency in handling the imaging probe, it is shown that varying the imaging plane to display various rows of microrobots is feasible. Acoustic stimulation parameters: $f = 101.0$ kHz and $V_{PP} = 27.3$ V.

**Movie S4.**
Depp tissue experiment: Five microrobots are visualized in real-time at an ultrasound phantom depth of 10 cm. Acoustic stimulation parameters: $f = 101.0$ kHz and $V_{PP} = 21.0$ V.

**Movie S5.**
Real-time motion experiment 1: The microrobot's spherical-like motion was recorded simultaneously using a CCD camera and the ultrasound imaging system in CFM-mode. Acoustic stimulation parameters: $f = 101.5$ kHz and $V_{PP} = 27.3$ V.

**Movie S6.**
Real-time motion experiment 2: The microrobot's linear-like motion was recorded simultaneously using a CCD camera and the ultrasound imaging system in CFM-mode. Acoustic stimulation parameters: $f = 101.5$ kHz and $V_{PP} = 27.3$ V.

**Movie S7.**
Drug-delivery approach: Acoustic microbubble streaming (1 μm-flow tracers) directed toward the wall can be utilized to perfuse dissolved drugs to the cavity's wall. Acoustic stimulation parameters: $f = 101.0$ kHz and $V_{PP} = 21.0$ V.
